# The non-canonical interaction between calmodulin and calcineurin contributes to the differential regulation of plant-derived calmodulins on calcineurin

**DOI:** 10.1101/2021.06.18.449055

**Authors:** Bin Sun, Xuan Fang, Christopher N. Johnson, Garrett Hauck, Jonathan P. Davis, Peter M. Kekenes-Huskey

## Abstract

Calmodulin (CaM) is an important Ca^2+^ signaling hub that regulates many protein signaling pathways. In recent years, several CaM homologs expressed in plants have been shown to regulate mammalian targets and they are attractive for gene therapy. However, the molecular basis of how the CaM homologs mutations impact target activation is unclear, which limits efforts to engineer their functional properties. To understand these mechanisms, we examined two CaM isoforms found in soybean plants that differentially regulate a mammalian target, calcineurin (CaN). These CaM isofroms, sCaM-1 and sCaM-4 share >90% and ~ 78% identity with the mammalian CaM (mCaM), respectively, activate CaN with comparable or reduced activity relative to mCaM. We used molecular simulations and experimental assays to probe whether calcium and protein-protein binding interactions are altered in plant CaMs relative to mCaM as a basis for differential CaN regulations. We found that the two sCaMs’ Ca^2+^-binding properties such as coordination and affinity are comparable to mCaM. Further, the binding of CaM to the CaM binding region (CaMBR) in CaN is also comparable among the three CaMs, as evidenced by calculated binding free energies and experimental measured EC_50_ [CaM]. However, mCaM and sCaM-1 exhibited stronger binding with a secondary region of CaN’s regulatory domain that is weakened for sCaM-4. This secondary interaction is likely to affect the turnover rate (*k_cat_*) of CaN based on our modeling of enzyme activity and is consistent with our experimental data. Together, our data show how plant-derived CaM variants can alter target activation through interactions beyond Ca^2+^-binding and canonical CaMBR binding, which may extend beyond the mammalian CaN target.

## 2 Introduction

Calmodulin (CaM) is a 16.7 kDa Ca^2+^ sensor ubiquitously expressed in eukaryotic cells [1] with an invariant sequence. CaM regulates at least 300 targets in cellular processes including muscle contraction, proliferation and immune response [2, 3]. CaM is also essential to plant physiology in response to environment stimuli [4]. In contrast to vertebrates, plants have different CaM isoforms [5, 6]. Two soybean CaM isoforms, sCaM-1 and sCaM-4, share ~90% and ~78% identity with mCaM, respectively, and have been extensively studied [7–9]. Interestingly, these two plant CaM variants show differential or even reciprocal regulation abilities on human protein targets [10, 11]. As an example, for calcineurin (CaN), a phosphatase, the sCaM-1 exhibits similar ability to activate the protein target as human CaM while sCaM-4 has significantly reduced ability [10]. CaN is a ubiquitously expressed serine/threonine phosphatase in all human tissues [12] and is regulated by CaM in a Ca^2+^-dependent manner. Disregulation of CaN is related to diseases such as autoimmunity [13], ventricular hypertrophy [14] and Alzheimer’s disease [15]. The positive correlation between sequence similarity and activation of CaN of these soybean CaMs with mCaM makes them good model systems to study the effects of CaM mutation on target activation.

CaM mutations found in human have been reported to disturb Ca^2+^-handling in cardiac cells [16]. To date the underlying molecular basis remains unclear. Several mechanisms may explain altered functions in CaM variants. CaM’s Ca^2+^binding property undoubtedly plays an important role in its target activation. It has been widely accepted that Ca^2+^binding promotes CaM’s N/C domains to expose the hydrophobic patch for target binding [17–19]. Although apo-CaM can bind targets [20], presence of Ca^2+^facilitates the formation of native complex [21], or even reverses CaM’s activating role [22]. Besides Ca^2+^-binding, the association between CaM and the CaM binding region (CaMBR) of targets is also important. A single mutation in the CaMBR region could abolish CaM binding [23]. Lastly, interactions between CaM and target region beyond the CaMBR can affect target regulation as well. For example, the ryanodine receptor (RYR) isoforms are differentially regulated by CaM, albeit bearing the same CaMBR region [24], suggesting that isoform-dependent region beyond CaMBR contributes to regulation. Another example is the activation of calcineurin (CaN) by CaM, in which a ‘distal helix’ region C-terminal to CaMBR interacts with CaM to fully remove the autoinhibitory domain of CaN [25]. Namely, studies [25–29] have provided a model that explains the molecular interactions between CaN and CaM that are vital to CaN’s complete activation (Fig. 1) [30]. In the model, Ca^2+^-saturated CaM binds the CaM binding region (CaMBR) (A391-R414) on the CaN regulatory domain, followed by a secondary interaction between CaM and a ‘distal helix’ (DH, K441-K466) that ultimately removes the auto-inhibitory domain (AID) from the CaN catalytic domain. These mechanisms could be utilized by CaM variants to alter target regulations.

**Figure 1:**
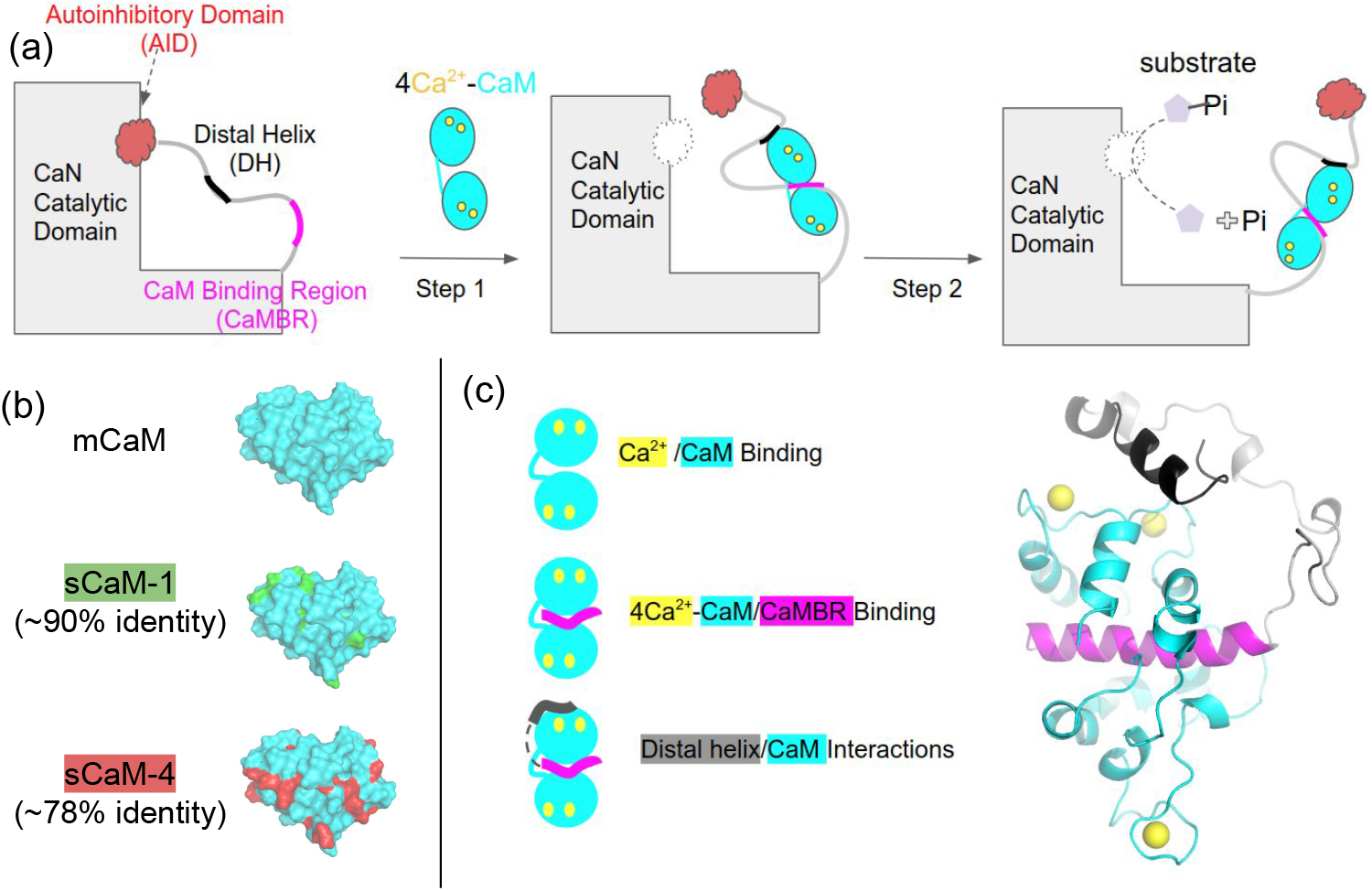
(a) Proposed CaN activation model by CaM [30]. Step 1, Ca^2+^-saturated CaM binds to the CaM binding region (CaMBR) and the distal helix (DH) motif of CaN, which displaces the autoinhibitory domain (AID) from its catalytic site. Step 2, Entry of substrate to CaN’s catalytic site enables dephosphorylation of substrate. (b) Two soybean CaM isoforms sCaM-1 and sCaM-4 share ~90% and ~78% sequence identity with mammalian CaM (mCaM). The unique residues are colored green and red for sCaM-1 and sCaM-4, respectively. These two sCaMs have differential regulations on human CaN. Detailed sequence comparison is in Fig. S1. (c) The three molecular interactions we examined in this study to explore the origins of the two sCaMs’ different activating abilities on human CaN.

In the present study, we therefore used CaN/CaM as a model system to explore the molecular basis of CaM target activation with different variants. We explored the origination of the two sCaM’s differentiated regulation abilities on human CaN. We hypothesised that the soybean CaM variants use one of these mechanisms to alter mammalian CaN activity: 1) Ca^2+^ binding properties to CaM, 2) CaM binding to CaMBR of CaN, and 3) region of the CaN regulatory domain distal to the CaMBR, the ‘distal helix’ motif. We found that the two sCaMs maintain comparable Ca^2+^ and CaMBR binding abilities to mCaM. However, the distal helix/CaM interaction patterns are different. This implicates the distal helix interaction as the cause for the different activation capabilities. We further pinpointed a hot-spot region on CaM that contributes most to the distal helix interaction difference between the two sCaMs and mCaM, This region lies in between residues N60-E87 and can be used in future CaM reengineering studies. Lastly, using a revised activation kinetic model, we found that CaM activates CaN by converting it to an activated form that exhibits altered kinetic properties. Altogether, our computational and experimental results showed that DH interactions are central to CaN activation by CaM.

## 3 Materials and methods

### 3.1 Calcineurin phosphatase assay using MUF-P substrate

The mCaM and sCaM-4 were overexpressed in E. coli and purified as described in [31] using phenyl sepharose. CaN was overexpressed from a pETagHisCN plasmid using E. coli BL21 (DE3) CodonPlus RIL cells. Cells were lysed by sonication, and CaN was purified using Ni-NTA and CaM-sepharose. Protein purity (>95%) was determined by Coomassie staining and SDS denaturing gel electrophoresis. Purified CaN fractions were pooled and dialyzed into 10 mM MOPS, 150 mM KNL, 1 mM TCEP at pH7.0. CaN and CaM concentrations were determined by UV-VIS spectroscopy. Experimental sample conditions were 2 mL of 150 CaN in assay buffer comprised of 200 mM MOPS, 150 mM KCl, 3 mM MgCl_2_, 2 mM EGTA at pH 7.0. The MUP concentrations was set to 0, 100, 200,400 and 600 *μ*M for substrate titrations of CaN and held at 100 *μ*M for EC_50_ experiments. Assay buffer EGTA concentration was calibrated based on a dose response of CaM tyrosine fluorescence (corresponding to CaM-C domain) against Ca^2+^. PerkinElmer spectrometer parameters were as follows: excitation 365 nm, emission 445 nm, data sampling interval 0.2 second, temperature = 20 °C. CaM titrations ranged from 5 nM to 3 *μ*M. Ca^2+^ titrations ranged pCa9 to pCa4. A minimum of 3 min. of raw fluorescence data were collected for each titration point and fit to a linear line using Kelidograph. The slope values were normalized and plotted against the titrant (CaM or Ca^2+^). Data was fit to a 4 parameter dose response curve using Prism. The resulting EC_50_ values were plotted in prism. Statistical significance was determine (95% confidence, p< 0.05) using an Anova test.

### 3.2 Kinetic Analyses

To investigate the activation effect of CaM on CaN, we first fit the Michaelis-Menten model to the kinetic data. It should be noted that due to the inner filter effect of MUF at high concentrations (data not shown), the kinetic assays were done at low concentrations. To estimate the kinetic parameters, the double reciprocal form of the model was fit to the data:

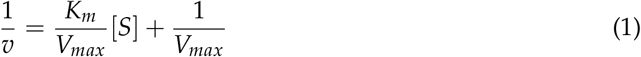

This Michaelis-Menten model is based on the assumption that activator binds to both the enzyme and the enzyme-substrate complex. However, in the activation of CaN by CaM, multiple molecular interactions are involved and thus a more sophisticated model is necessary.

To gain further insight into the activation mechanism of CaM and compare the kinetic properties of different CaM isoforms, we proposed a revised activation model for the data (Fig. 2). In this revised model, we assume Michaelis-Menten mechanism for isolated CaN. However, upon binding to CaM, CaN is then converted into the activated form that has different kinetic properties as specified by the *α* and *β* factors. The fitting of the revised model was done in two steps. For the CaM free data set, the *K_m_* and *V_max_* was determined by fitting the double reciprocal form of the Michaelis-Menten model to this data set. We then assumed that mCaM and the soybean isoforms have the same *K_a_* (0.038 ±0.0048 *μ*M), which we validated in this study via experimental assays. Together with the *K_m_* and *V_max_* determined from the previous step, the reciprocal form of the revised model (Eq. 2) was fit to the CaM and sCaM data sets with *K_m_*, *V_max_*, and *K_a_* as shared and fixed parameters (*K_m_* and *V_max_* are intrinsic properties of CaN) while *α* and *β* were used as free parameters.

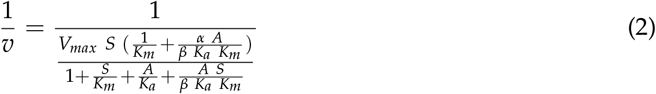

where A is CaM, *K_a_* is the binding constant of CaM to CaN, *α* is scales of *V_max_* and *β* is scales of *K_m_*.

**Figure 2:**
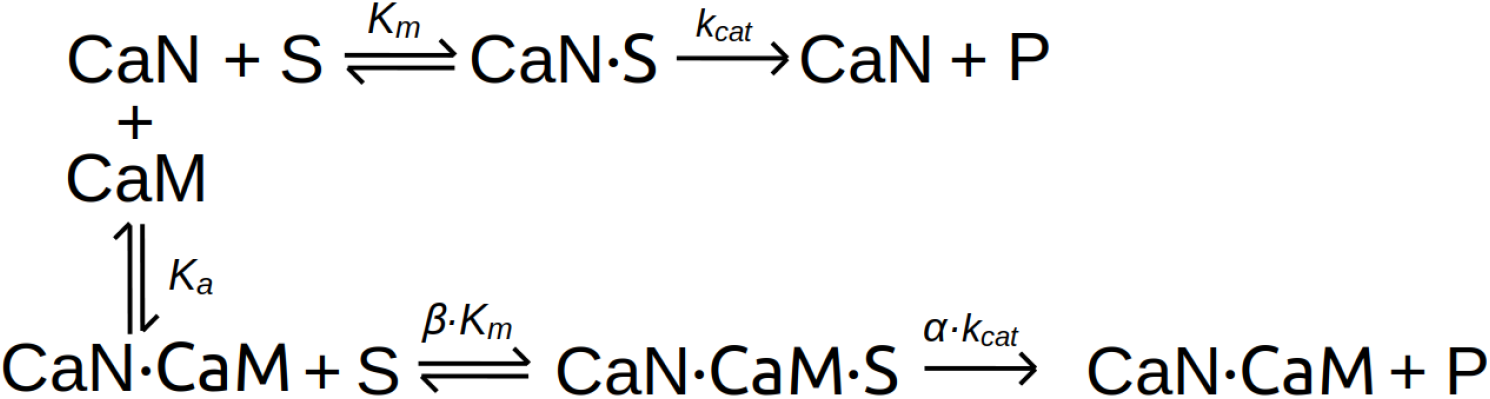
A kinetic model that considers multiple molecular interactions involved in CaN’s enzymatic activity after being activated by CaM.

### 3.3 Rosetta Comparative Modeling of sCaM-1 and sCaM-4 structures

The available structural information for sCaM-1 and sCaM-4 deposited in PDB databank are for isolated N- or C-domain structures [11] (PDB ID: 2RO8/2RO9 for sCaM-1 and 2ROA/2ROB for sCaM-4, respectively). Although there is one complete sCaM-4 structure in complex with vacuolar calcium ATPase BCA1 peptide deposited with PDB ID 2L1W [7], the binding mode of target peptide is significantly different relative to the mammalian CaM/CaMBR structure. For the sCaM structure, the N- and C-domain binds to the two ends of peptide, as opposed to the conventional ‘wrap round’ manner in which N-/C-domain collapse around the middle part of target peptide. Therefore, we built complete sCaM-1 and sCaM-4 structures that bind CaMBR in the same manner as mammalian CaM. The complete structures of sCaM-1 and sCaM-4 were built using the Rosetta comparative modeling approach [32]. For each model, the mammalian CaM/CaMBR complex crystal structure (PDB ID 4Q5U) as well as the N-/C-domain structures were used as templates (the weights of each template are 1.0, 0.8 and 0.64, which are the default values in the RosettaCM hybridize mover). The hybridize function of Rosetta was used and the specific protocol and flags are given in Sect. S2.1. For each soybean CaM, the highest ranked structure generated by Rosetta comparative modeling was subject to 3 × 2 *μ*s all-atom MD simulation refinement.

### 3.4 All-atom MD simulations to refine the modeled sCaM structures

All-atom MD simulation was performed to refine the Rosetta modelled soybean CaM structures. The ff14SB [33] force field was used for protein atoms. The system was solvated in TIP3P [34] waterbox with the distance between protein to waterbox wall being 14 Å. K^+^ and Cl^-^ ions were added to maintain 0.15 M salt concentration. The system was subjected to an energy minimization, for which all atoms except hydrogens, water and KCl ions were constrained by the ibelly functionality. The cutoff value for non-bond interactions was set to 10 Å. A 2 fs timestep was chosen, as SHAKE [35] constraints were applied on bonds involving hydrogen atoms. Two heating procedures were performed to heat the system from 0 to 300 K using the Amber18 sander.mpi engine [36]. In the first heating stage, the *ibelly* function was used to keep all system except the water box and KCl ions fixed. The water box was heated to 300 K over a 100 ps interval under the NVT ensemble. For the second heating stage, the entire system was heated from 0 to 300 K over 500 ps under the NPT ensemble, for which the backbone atoms were constrained by an harmonic potential (force constants of 3 kcal mol^−1^ Å^−2^). Thereafter, an additional 1 ns equilibrium stage was conducted at 300 K under the same constraints, but with a reduced force constant of 1 kcal mol^−1^ Å^−2^. The Langevin dynamics was used during the simulation. These equilibrium simulations were followed by triplicate 2 *μ*s production-level molecular dynamics (MD) simulations. The simulation length is sufficient to refine the Rosetta modelled structure and give reliable sCaM structure. This is based on the reports that the timescale of conformational fluctuation in the MD refinement of modeled structure is approximate 1 *μ*s for systems with mean residue number around 120 [37], which is approximate the length of CaM.

### 3.5 Docking distal helix structure to CaMs

#### 3.5.1 Replica exchange molecular dynamics (REMD) simulations of isolated distal helix region

REMD was performed on the distal helix fragment (residues K441-K466 of CaN, amino acid sequence is shown in Fig. S6) to sample representative conformations of the isolated DH. 12 replicas of the system were constructed and the temperature range of the REMD was 270 to 454.72 K, which was calculated based on the protocol proposed in [38]. The temperature range ensures that the exchange probability between neighbouring replicas is approximately 0.4 as proposed in [39]. The system was parameterized with ff99SBildn force field [40] coupled with an generalized Born implicit solvent model. The detailed REMD simulation procedure has been described in [30]. The production run of the REMD was 100 ns and clustering analysis were performed on the trajectory from 297 K. Representative structure of the most populated cluster was selected for docking to CaM structures. The selected DH conformation is shown in Fig. S6.

#### 3.5.2 Docking DH to CaM surface and extensive MD refinement

The representative structure of the DH generated by REMD simulations was docked to surface of sCaM-1, sCaM-4 and mCaM via the ZDOCK webserver [41]. The CaM structures that were used for the docking are from the MD refinements of Rosetta modeled structures. Specifically, clustering analysis were first performed on the MD trajectories and the representative structure of the most populated cluster was selected. For mCaM, one representative structure was chosen and for the two sCaMs, two representative structures were chosen for each case. The procedure of clustering analysis and identifying the two representative structures is detailed in Sect. S2.3. After preparing the DH and CaM structures, the DH was docked to CaM following the docking procedure in [30]. Four putative binding sites were pre-defined on CaMs that locate closely to the grooves formed by helix bundles, as these places are more likely to accommodate helix-helix interactions [42]. The selected four putative sites covered most of the CaM’s solvent-accessible surface. The residues in Table S2 from each site on CaM were specified as contact residues during docking to ensure the initially generated distal helix poses are near the putative binding sites. The detailed docking procedure was reported in [30].

The highest-scored CaM/DH interaction pose at each site was selected for further MD refinement with triplicate 1.2 *μ*s runs. The detailed procedure of docking and MD refinement has been reported in [30]. In summary, 72 *μ*s refinement MD simulations were performed with 28.8 *μ*s for each sCaM-1 and sCaM-4 and 14.4 *μ*s for mammalian CaM. All molecular dynamics simulations were performed via Amber16 package [43] and are summarized in Table S1.

### 3.6 Analyses Methods

Clustering analysis, root mean squared deviations (RMSD)/root mean squared fluctuations (RMSF) calculations, hydrogen bonds, and secondary structure analysis were performed via cpptraj [44]. The reference structure used for these analyses was the CaM/CaMBR crystal structure (PDB ID: 4Q5U [27]). Secondary structure for each residue was calculated using cpptraj with the Define Secondary Structure of Proteins (DSSP) algorithm [45].

#### 3.6.1 Projection of MD sampled conformations onto 2D plane using the sketch-map dimensionality reduction method

Sketch-map (https://github.com/cosmo-epfl/sketchmap) is a non-linear dimensionality reduction method that we used to visualize CaM/DH complex conformation space sampled in the MD simulations [46, 47]. Each frame from the MD trajectories was represented by a point in the high-dimensional space 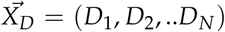 and *D*_1_, *D*_2_…*D_N_* are the minimums distance between *C_α_* atoms of CaMBR and distal helix. In the present study, N was set as 32: *D*_1_ to *D*_14_ represents the minimum distance of the 14 residues in CaMBR (E394-A407) to the distal helix. Namely, for residue *i*, we first calculated its distance to each residue in the distal helix and assign the smallest value to *D_i_*. And *D*_15_ to *D*_32_ are the reverse minimum distances, namely, the minimum distance of each residue in the distal helix (S438-D455) to CaMBR residues. This definition of high-dimensional points ensures that the relative conformation information between distal helix to CaM/CaMBR is entirely encoded in the coordinates of high-dimensional points, as illustrated in [48]. These high-dimensional points were projected onto a 2D plane as described in [48] to generate the sketch-map representation of MD sampling.

## 4 Results and Discussion

### 4.1 Experimental assays revealed that sCaM-1 and mCaM exhibited stronger activation of CaN than sCaM-4

We anticipated that sCaM-1 would more strongly activate human CaN relative to sCaM-4, based on the form’s higher sequence similarity to mCaM. Indeed, a study by Cho *etal.* demonstrated that sCaM-1 exhibited a stronger activation of CaN relative to sCaM-4, based on their measurements of CaN activity as a function of CaM with 100 *μ*M MUF-P as a substrate [10]. To investigate the kinetic mechanisms by which the CaM variants differentially activate CaN, we performed pMUF substrate titrations of CaN alone as well as CaN with different CaM isoforms. As shown in Fig. 4, the presence of CaM (regardless of the isoform relative to isolated CaN) significantly increased the activity of CaN. Consistent with Cho *etal*. observations, sCaM-1 enhanced CaN’s activity to a similar extent as mCaM while sCaM-4 did so to a much lesser extent.

**Figure 3:**
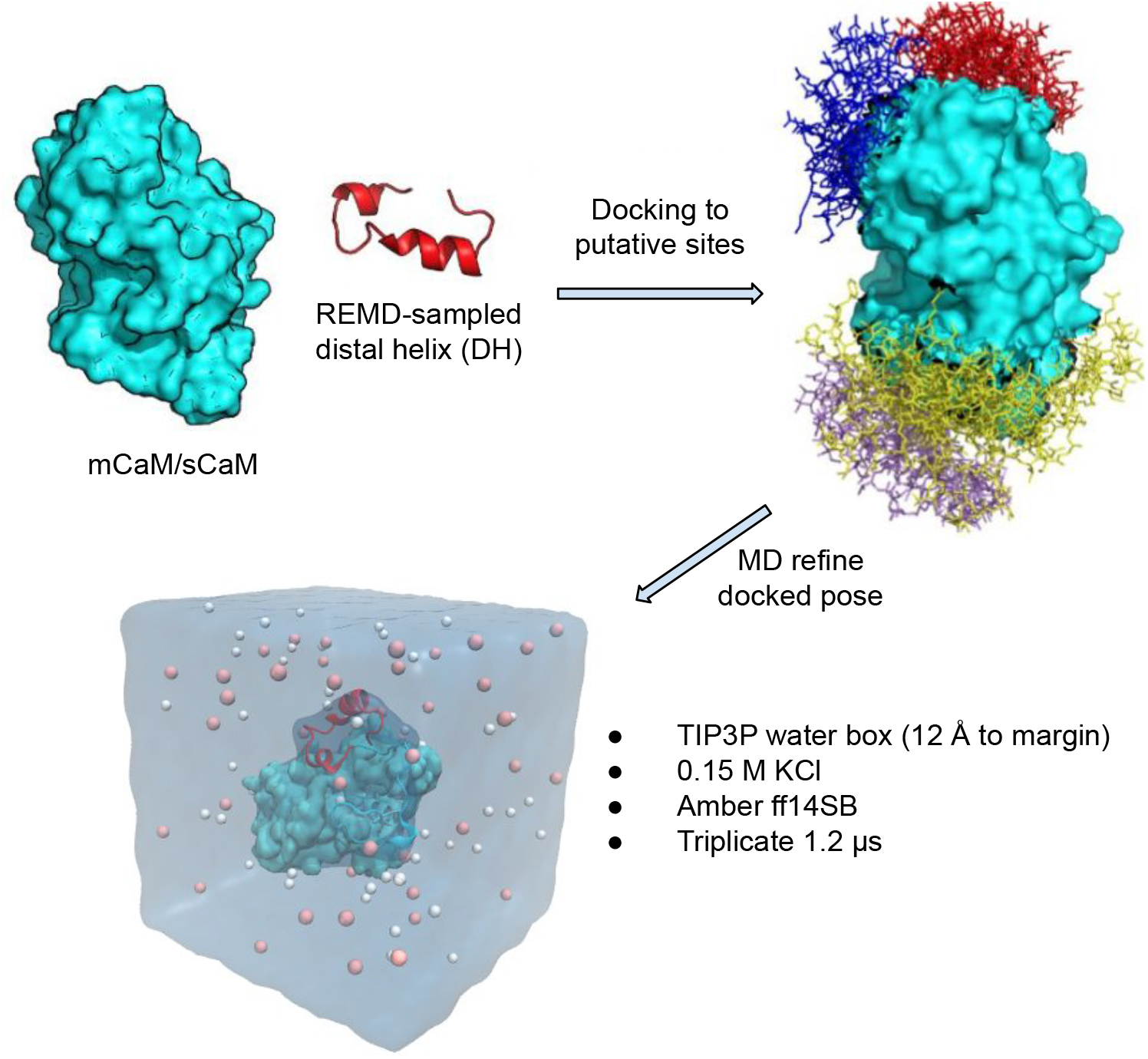
Scheme of docking the REMD-sampled distal helix (DH) conformation to CaM’s surface via the ZDOCK webserver [41]. For each CaM case, four putative binding sites on CaM that are localized to the grooves between helices were pre-defined as contact site during the docking. The putative site residues of CaMs are listed in Table S2. For each site, the highest-scored pose was further subjected to all-atom MD refinement.

**Figure 4:**
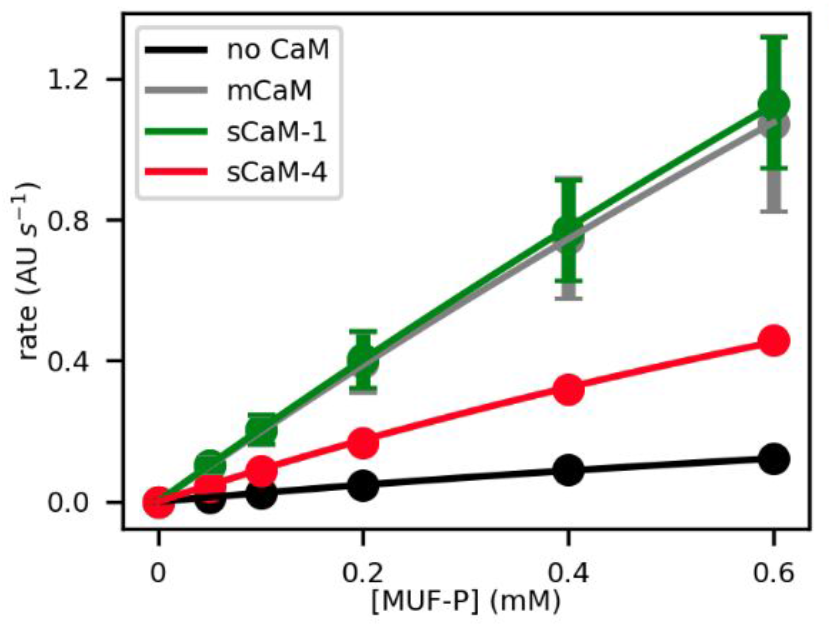
Experimental CaN activity. Substrate titrations of CaN alone as well as CaN with different CaM isoforms.

### 4.2 The different activation effects between sCaM-4 and mCaM/sCaM-1 are not caused by their Ca^2+^ binding properties and their CaMBR binding properties

We next sought to determine the molecular mechanisms behind these experimental trends *via* computational modelings, which included homology modeling, protein-protein docking and molecular dynamics simulations. As shown in Fig. 1a, the previous model for CaM/CaN activation includes 1) Ca^2+^ binding to isolated CaM. 2) binding of Ca^2+^-saturated CaM to the CaM binding region (CaMBR) of CaN and 3) the interaction of the distal helix (DH) motif from from CaN to CaM. We examined theses interactions for the mCaM and sCaM isoforms in the activation of CaN.

Because the complete sCaM-1/4 structures complexed with CaMBR of CaN are not yet available, we performed homology modeling followed by extensive all-atom MD refinement to obtain structural models. We show in Sect. S2.2 that the modelled sCaM/CaMBR complex structures were reasonable as supported by converged RMSD values over the *μ*s MD time-scale, low fluctuations of backbone atoms (<2 Å RMSF for most part) and a more compact structure of sCaM-4 that was consistent with experimental observations (Fig. S3). Based on the modelled structures, we examined key molecular interactions that may contribute to the sCaM isoforms’ differential activation of mammalian CaN.

#### 4.2.1 Ca^2+^ binding property

Ca^2+^ plays a vital role in CaM’s target regulation [49]. In this section we examined if the Ca^2+^-binding properties were similar among mCaM and the sCaM isofroms. CaM has four EF hands, two in each N-/C-domain. MD simulations indicated that the EF-hands in the modelled sCaM structures were stable as evidenced by the ~3 Å RMSF values (Fig. S3). Not surprisingly, mCaM had the smallest RMSF of ~1 Å at EF-hands. sCaM-1 had the largest RMSF among the three CaMs because the anti-parallel *β*-sheet between EF-hands was disrupted. We anticipated this was due to a structural organization to maintain Ca^2+^ affinity, similar to *β*-sheet reductions observed in parvalbumins upon Ca^2+^ binding [50].

Nonetheless, sCaM-1 maintained the same Ca^2+^ coordination number as the other two CaMs. Fig. S4(a,b) reports the integrated radial distribution functions (IRDF) of amino acid oxygens and water oxygens around Ca^2+^. In previous studies, we have shown intrinsic Ca^2+^affinities in EF-hands tend to correlate with oxygen RDFs [50, 51]. In all CaM isoforms, the IRDF curves reached a plateau around 2.3 Å, which indicates the average distance between Ca^2+^ and the coordinating oxygens RDFs were indistinguishable across all cases, which suggests similar Ca^2+^ binding affinities. Interestingly, as shown in Fig. 5a, the number of amino acid oxygens and water oxygens varied somewhat: sCaM-1 had the lowest amino acid oxygen number around 5.5 for Ca^2+^ at the firs EF-hand of N-domain and second EF-hand of C-domain but was compensated for by the highest water oxygen numbers around 2. In addition, the Ca^2+^s bound at the C-domain of the two sCaMs had more water oxygens compared with mCaM. Nevertheless, the total Ca^2+^ coordination numbers were comparable, all were in the range of seven to eight oxygens (Fig. 5a), in agreement with the Ca^2+^ coordination number measured from other Ca^2+^-binding proteins [30, 51–54] and in aqueous solution [55]. The one oxygen variation should have negligible impact on Ca^2+^ binding free energy in the EF-hand site as we demonstrated previously difference of ±1 oxygen do not appreciably impact Ca^2+^ binding free energy in the EF-hand sites [50]. Since changes in Ca^2+^ affinity can enlist conformational changes outside of the EF-hands [50, 56], we experimentally measured the EC_50_ [Ca^2+^] of mCaM and sCaM-4 in activating CaN (Fig. 6(b-c)). This was obtained by measuring CaN activity at varying Ca^2+^ concentrations with saturated CaM. The mean values of the EC_50_ for mCaM and sCaM-4 were 0.68 and 0.49 *μ*M, respectively. Converting this EC_50_ to energy via – *RT* ln *EC*_50_, which implies 0.19 kcal/mol energy difference between mCaM and sCaM-4 at room temperature, and thus Ca^2+^ binding properties were comparable. These data together showed that the three CaM isoforms maintained comparable Ca^2+^ binding properties, which suggested the differential CaN activation ability was not caused by Ca^2+^ binding.

**Figure 5:**
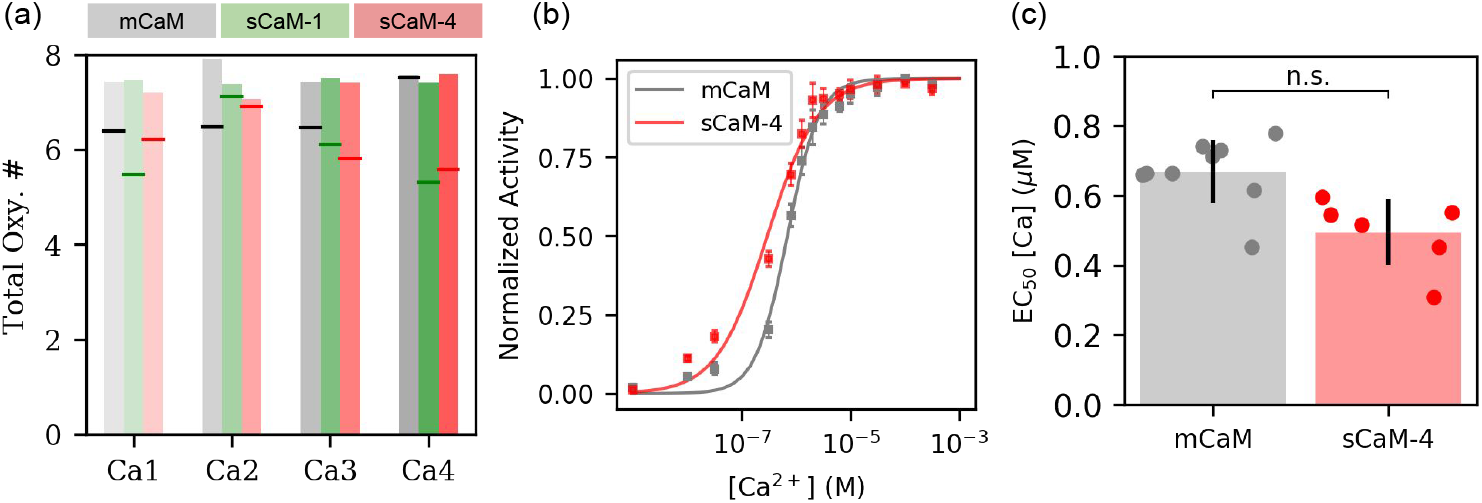
Ca^2+^ binding properties. (a) The total number of coordinating oxygens (amino acid oxygen plus water oxygen) around Ca^2+^. The two Ca^2+^s at the N-domain EF-hands were indexed as “Ca1” and “Ca2”, and the Ca^2+^s at C-domain were indexed as “Ca3” and “Ca4”. The horizontal lines within each bar indicate the number of amino acid oxygens. (b) Dependence of CaN activity on Ca^2+^under saturating CaM condition. (c) Experimental EC_50_ [Ca^2+^] for mCaM and sCaM-4. n.s. p >0.05.

**Figure 6:**
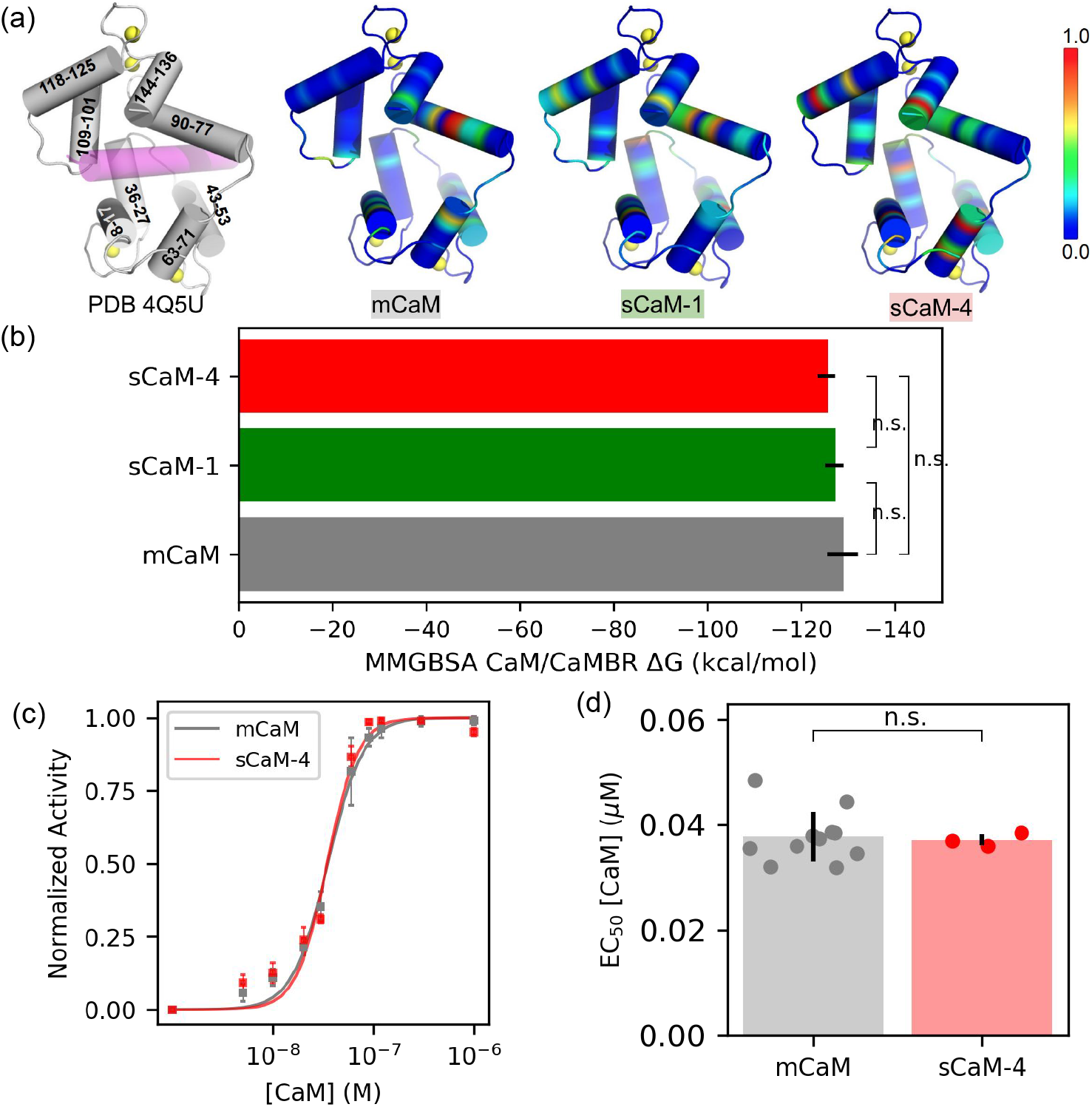
CaMBR binding properties. (a) Normalized contact probability (with respect to the residue pair that has the most accumulated contacts from the MD sampling) of CaMBR with CaM projected on mCaM/CaMBR complex structure PDB 4Q5U. The residues numbers comprising the helices are labeled. (b) MM-GBSA calculated binding free energy between CaMBR and CaM using MD trajectories. The difference between the three CaMs were non-significant as determined by student’s t-test. (c) Dependence of CaN activity on CaM under saturating Ca^2+^ condition. (d) Experimental EC_50_ [CaM] for mCaM and sCaM-4.

#### 4.2.2 CaMBR binding property

We next examined if the binding between the CaMBR and CaM differed for the three CaM isoforms, as a basis for their different activities. We reported in Fig. 6a the contacts between CaMBR and CaM. Overall, the CaMBR/CaM interaction pattern shown in mCaM were largely maintained by the two sCaMs, especially the contacts from CaM helices formed by residues Q8-S17, T43-N53, K77-R90. In the simulations of all CaMs, the CaMBR exhibited small fluctuations with RMSF < 2 Å (Fig. S3b), which suggested that the CaMBR was highly stable when bound to three CaMs. The largest difference was observed in the C-domain: both sCaMs exhibited contacts with the CaMBR via two helices through residues S101-M109 and D118-I125, while mCaM had negligible contacts with CaMBR at these two helices. However, the MMGBSA-calculated binding free energies between the CaMBR and CaM (Fig. 6b) were comparable across the three CaMs and were highly favorable: −128.8 ±3.3, −127.1 ±2.0, −125.4 ±1.9 kcal/mol for mCaM, sCaM-1 and sCaM-4, respectively. To verify, we measured the EC_50_ [CaM] for mCaM and sCaM-4 in activating CaN. This was obtained by measuring CaN activity with varying CaM concentrations under saturating Ca^2+^. As shown in Fig. 6c, the dependence curves of CaN activity on mCaM and sCaM-4 were indistinguishable and the calculated EC_50_ [CaM] for mCaM and sCaM-4 were equal (~0.038 *μ*M, Fig. 6d). Although EC_50_ is not merely the CaM binding affinity towards CaN, CaM and CaMBR binding affinity contributes large part to the measured *E*C_50_. This is because CaMBR region is universal in CaM-targets and this region is the primary CaM recognition site when activating targets [2]. Thus, the almost identical EC_50_ values for mCaM and sCaM-4 suggest that these two CaMs are highly likely to have the same CaMBR binding affinities. Additionally, it has been shown that the two sCaM’s bind to the target with comparable affinities and are also comparable to other mCaM/targets binding affinities [11,20], which support our interpretation of EC_50_ [CaM] data. The two sCaM did not show significant difference in binding CaMBR when compared with mCaM, based on 1) Most of the contacts shown in CaMBR/mCaM binding are maintained in the two sCaMs. 2) The MMGBSA-calculated CaMBR/CaM binding free energies are comparable. Therefore the different CaN activation ability of sCaMs is not likely due to CaMBR binding.

### 4.3 mCaM/sCaM-1 exhibited stronger binding with distal helix than sCaM-4

We next examined if CaM/distal helix binding differed among the three CaMs as the final potential mechanism for differential CaN activation (Fig. 1c). To analyse the CaM/distal helix binding models, we first performed 100 ns REMD simulation of the isolated distal helix to sample the representative distal helix conformation as initially described in [30] (see Fig. S6 for REMD results). The representative distal helix was then docked to CaM isoforms via protein-protein docking, followed by extensive all-atom MD simulations. To visualize the CaM/distal helix conformational space, the MD trajectories were projected onto a 2D plane via the Sketch-map method [46], using the distal helix to CaMBR distance as the coordinate basis (Fig. 7a).

**Figure 7:**
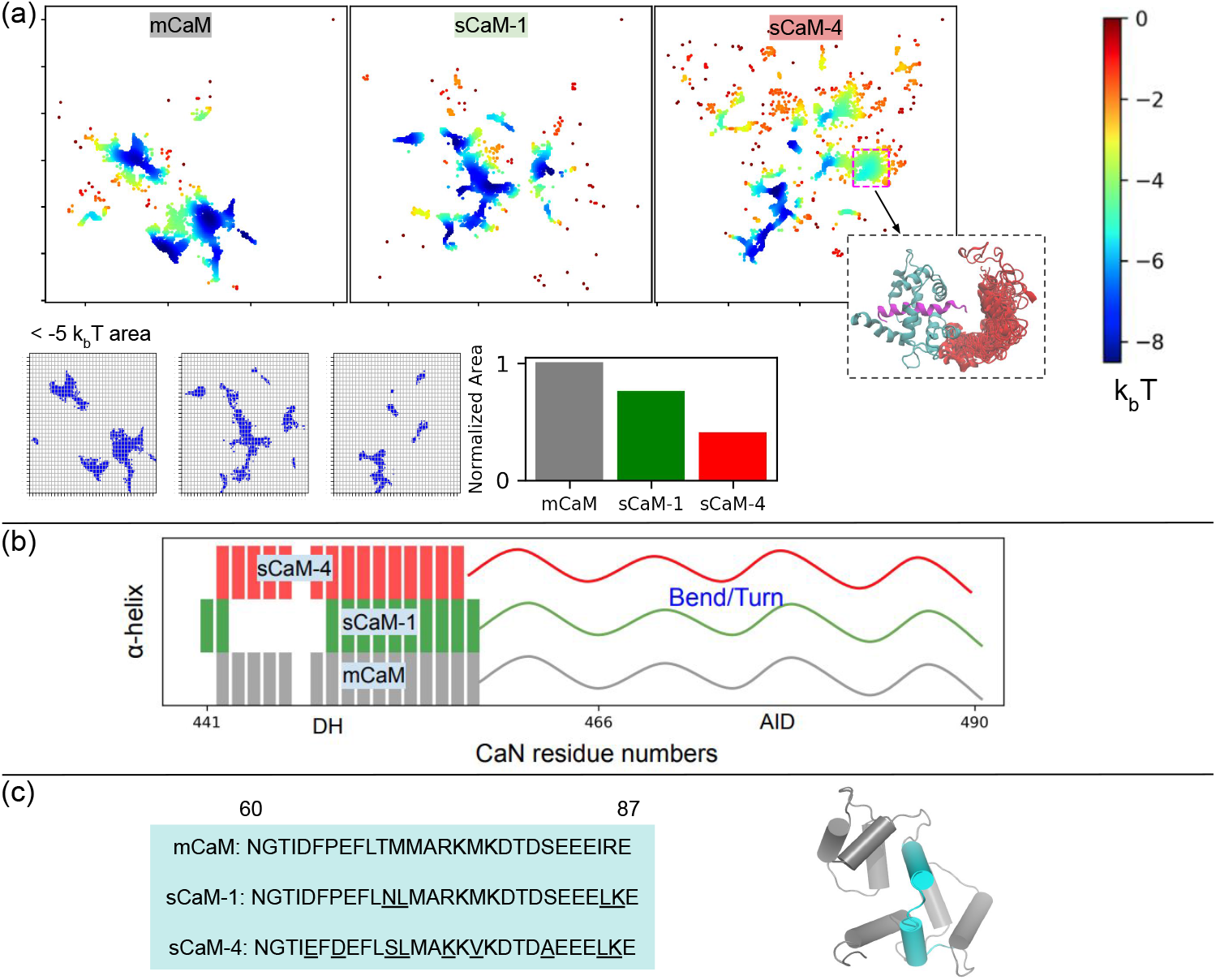
(a) Poses of the distal helix (DH) interacting with CaM. The poses are colored based on their energies using −*k_b_T* ln(*ρ*/*ρ_o_*) where *k_b_* is Boltzmann constant, *T* is temperature and *ρ* is the point density after projection using Sketch-Map dimensionality reduction method [46]. *ρ_o_* is density of the peripheral points in sCaM-4 (dark red points). Region that are below −5 *k_b_T* are used as a reference for stable binding. In sCaM-4, representative structures from unstable region (> −5 *k_b_T*) are shown with DH colored red. (b) Overall *α*-helicity of residues comprising the DH and AID of CaN from MD simulations. A bar depicts a residue that has *α*-helix as dominant secondary structure, which was calculated via the CPPTRJ program [44]. (c) Sequence comparison of residues N60-E87 of two sCaMs with mCaM. The region contributes most to the DH/CaM contact map difference between mCaM/sCaM-1 and sCaM-4 shown in Fig. S7. Non-conserved residues are underlined. The location of this sequence region is highlighted as cyan in the mCaM crystal structure PDB ID: 4Q5U.

The densities (*ρ*) of the projected points reflected the distribution of DH poses on CaM’s surface. For easier interpretation, we converted the densities to free energies using −*k_b_T* ln(*ρ*/*ρ_o_*) where *ρ_o_* is the density corresponding to the peripheral points in sCaM-4. As shown in Fig. 7a, the free energy minimum of DH/CaM interaction were all about 8.5 kcal/mol for all three cases, which suggested that there were thermodynamic-favourable binding sites for the distal helix on all CaM isoforms. This is consistent with our experimental data demonstrating that all CaM isoforms are capable of activating CaN. Using < −5*k_b_T* as an arbitrary criterion to define thermodynamic favourable area, mCaM had the largest area while sCaM-1 and sCaM-4 had 75% and 40% of mCaM, respectively (Fig. 7b). Visual inspection showed that high energy conformations of DH from sCaM-4 were more spread instead of being concentrated on CaM The larger interacting area provided by sCaM-1 for the distal helix interaction is consistent with sCaM-1’s complete activation of CaN versus partial activation by sCaM-4 (see Fig. 4).

We next performed contact map analyses to identify the key CaM residues that contributed to CaM/DH interactions among these three CaMs. We found that mCaM had the strongest contact in terms of number of contacting residues as well as contact persistence measured by the accumulated contacts (Fig. S7). sCaM-1 had comparable contact persistence with sCaM-4 but with more pairs of contacts. The notable difference was that mCaM residues (T62, I63 and D64) and sCaM-1 resides (D64, F65, P66 and F68) had more contacts with DH than sCaM-4 (Fig. S7). These residues locate closely to the linker of CaM that connects the N- and C-domains. In this region, sCaM-1 has six different residues relative to mCaM while sCaM-4 has ten (Fig. 7c). The larger sequence difference of sCaM-4 relative to mCaM/sCaM-1 is likely to weaken the distal helix interaction. sCaM-4 exhibited weaker CaM/DH interaction than mCaM and sCaM-1, which likely contributes to sCaM-4’s reduced CaN activation ability.

**Table 1:**
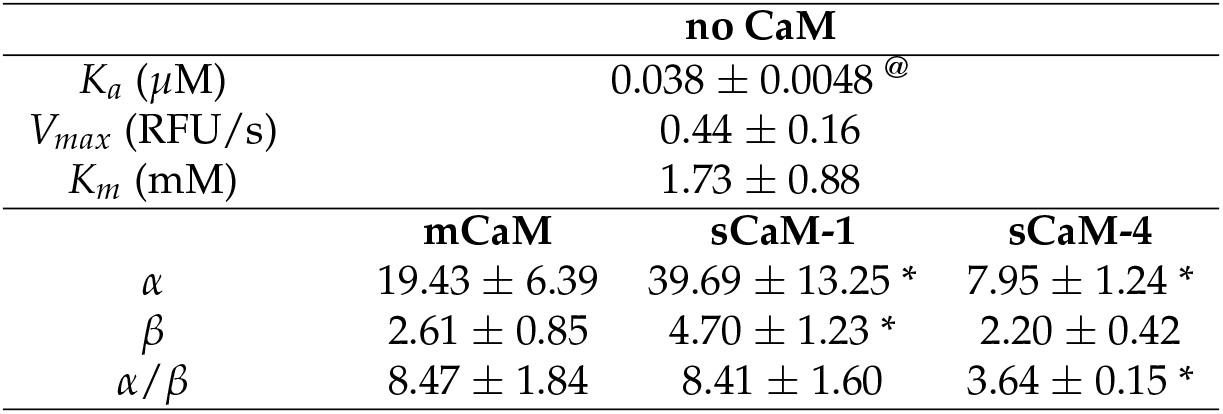
Kinetic parameters obtained by fitting model in Fig. 2 to experimental data. Student’s t test was used to determine the significance of difference between mCaM and the soybean isoforms. * p < 0.05. @ obtained from EC_50_[CaM] data in Fig. 6

Higher helical content of DH may facilitate the AID displacement via shortening the distance between DH and AID. We calculated this propensity and determined that the overall *α*-helical residues in DH were comparable for these three CaM (Fig. 7b). The distal helix in all cases adopted *α*-helix structure at the C-termini. When interacting with CaM, the number of *α*-helical residues were comparable for these three CaM, with 16, 12, and 15 residues for mCaM, sCaM-4 and sCaM-4, respectively. Hence, the CaMs don’t appear to impact distal helix upon activating CaN. This suggests distal helix helicity may be a precursor to activity showed by these CaM isoforms.

### 4.4 Kinetic fitting suggested that stronger CaM/DH led to larger apparent *k_cat_* of CaN

To determine how CaM/CaN interactions impact CaN’s kinetics, beyond standard molecular modeling, we fit the kinetic data shown in Fig. 4 to the kinetic model in Fig. 2. This model includes additional CaM/CaN interactions studied via simulations. As shown in Fig. 1a, multiple interactions such as CaM/CaMBR interaction, CaM/DH interaction and activated CaN/substrate interactions are involved in the dephosphorylation process. There are three assumptions in the proposed model: 1) CaM binds only to the enzyme but not the enzyme-substrate complex, in accordance with the mechanism of CaN activation by CaM (Fig. 2). 2) The three CaM isoforms share the same *K_a_* as indicated by the EC_50_[CaM] data where mCaM and sCaM-4 showed nearly identical EC_50_[CaM] (Fig. 6). 3) Once bound by CaM and activated, CaN exhibits modified kinetic properties described by *α* and *β* factors which rescale k_cat_ and *K_m_*, respectively.

Firstly, fitting yielded an *α* value much greater than 1 for all CaM cases, indicating that CaM significantly enhanced the catalysis of CaN (*k_cat_*). All CaM isoforms also exhibited *β* > 1, indicating a modified *K_m_*. However, it should be noted that interpreting *K_m_* as the substrate affinity is not appropriate here for the following reasons, which are: Suppose the binding of S to CaN can be described by *k_on_* and *k_off_*, the on and off rates of S to CaN. 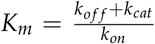 therefore approximates 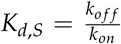 if *k_cat_* ≪ *k_on_*, *k_off_*. In this case, a change in *α* is independent of a change in *β*. However, as our kinetic data revealed, both *α* and *β* increased, implying a possible correlation between these two parameters. Consequently, interpretation of *K_m_* as the affinity for substrate is nuanced.

To examine the impact of different CaMs on CaN kinetics, we computed *α*/*β* as a measure of catalytic efficiency. Both mCaM and sCaM-1 showed nearly identical *α*/*β* and are about twice that of sCaM-4. This indicates that upon CaM binding, CaN is converted into an more activated form (*i.e*. AID-removed CaN), in contrast to its autoinhibited form. And more importantly, this activated form possesses enhanced catalytic capability. In the previous sections, both experimental and computational data demonstrated that the Ca^2+^ and CaMBR binding did not contribute much to the different activation effects of CaM isoforms. Meanwhile, the three isoforms exhibited differences in CaM/DH interaction. Hence the difference in catalytic efficiency could be attributed to the difference in DH binding. Kinetic analyses suggested that CaM increased the turn over rate. CaN activated by mCaM and sCaM-1 had significantly larger turn over rate (*k_cat_*) than that of sCaM-4.

To test this hypothesis, using a polymer model, we showed in Sect. S2.5 that the CaM/DH interaction is positively correlated with the effective AID concentration ([AID]_eff_), a quantitative measure of the extent of CaN being autoinhibited. In the case of mCaM and sCaM-1, the CaM/DH interaction led to lower [AID]_eff_ than that of sCaM-4. Hence, it is possible that the DH/CaM binding in mCaM and sCaM-1 pulled the CaN’s AID away from its catalytic site to a greater extent through the interactions with DH, resulting in a CaN form that showed higher catalytic efficiency (*α*/*β*). Further studies will be needed to delineate other potential mechanisms, including interactions of CaM with the well folded catalytic domain.

## 5 Conclusions

CaM is an important Ca^2+^ sensor that translates Ca^2+^ signaling to various protein-protein interactions. It was long believed that CaM has an invariant amino acid sequence in vertebrates [57] and has only one amino acid difference to several insect CaM sequences [58]. However, recent studies have revealed that CaM mutations in human populations have so far been related to cardiac diseases [31, 59], likely through disturbing Ca^2+^-handling in cardiac cells [16]. The molecular mechanisms underlying calmodulinopathy are still emerging [60]. Understanding how plant-derived CaM isoforms affect the activation of mammalian targets helps to explain the altered functions of CaM variants found in human. In the present study, we examined the molecular basis of two soybean CaM isoforms that differentially activate human CaN. Two soybean CaMs, sCaM-4 and sCaM-1, share moderate to high sequence identity to human CaM (mCaM), but exhibit differential regulation of human CaN: sCaM-1 was comparable with mCaM and sCaM-4 was significantly reduced. Based on a recent CaM/CaN interaction model [30], we investigated three molecular interactions that may affect CaM’s activation on CaN.1) Ca^2+^ binding to CaM.2) CaM binding to the CaM binding region (CaMBR) of CaN and 3) CaM binding to the distal helix (DH) motif of CaN. By using molecular dynamics simulations and experimental measurements, we excluded Ca^2+^ binding and CaMBR binding as factors that differentiates the sCaMs. Instead, it is the CaM/DH interaction that strongly differentiates sCaM-4 versus mCaM/sCaM-1. Namely, mCaM and sCaM-1 both exhibited stronger interactions with distal helix than sCaM-4. Kinetic modeling revealed that the prime effect of the two sCaMs was to form CaN/CaM complex with different turnover (*k_cat_*) rates, which is likely resulted from altered CaM/DH interactions. Using a polymer model, we found that the CaM/DH interaction correlates with the effective [AID] concentration around the catalytic site of CaN, which could impact the turnover rate. Our study shed insights into how CaM mutation may affect target activation through interaction beyond the canonical CaMBR binding and Ca^2+^ binding, which may extend to other CaM regulated targets such as CaM protein kinase II (CaMKII [61]), CaM kinase kinase (CaMKK [62]), myosin light chain kinase (MLCK [63] and death-associated protein kinase (DAPK) [64]) that utilize autoinhibitory domains.

## 6 Acknowledgements

Research reported in this publication was supported by the Maximizing Investigators’ Research Award (MIRA) (R35) from the National Institute of General Medical Sciences (NIGMS) of the National Institutes of Health (NIH) under grant number R35GM124977 (PKH). This work was also supported by American Heart Association under grant number 20CDA35310757 (CNJ) and NIH under grant 5R01HL138579-04 (JPD). This work used the Extreme Science and Engineering Discovery Environment (XSEDE) [65], which is supported by National Science Foundation under grant number ACI-1548562.

## S1 Supplementary Information (SI)

**Table S1:**
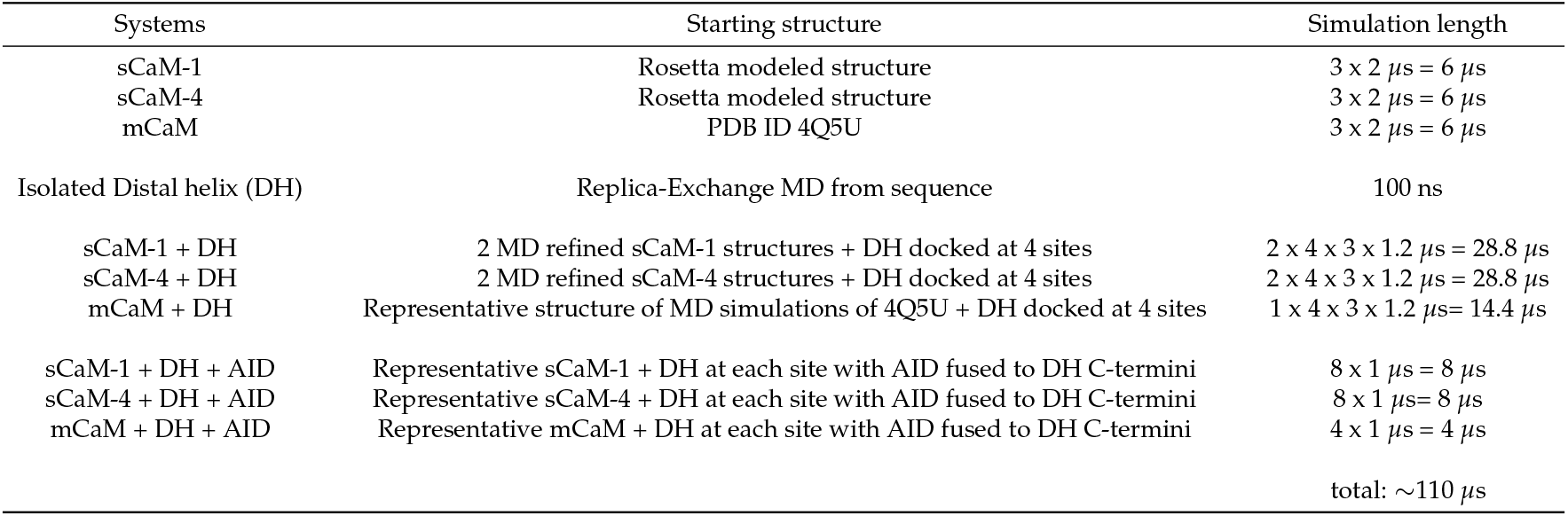
All molecular dynamics simulations performed in present study

| Systems | Starting structure | Simulation length |
| --- | --- | --- |
| sCaM-1 | Rosetta modeled structure | $3 \times 2 \mu s = 6 \mu s$ |
| sCaM-4 | Rosetta modeled structure | $3 \times 2 \mu s = 6 \mu s$ |
| mCaM | PDB ID 4Q5U | $3 \times 2 \mu s = 6 \mu s$ |
| Isolated Distal helix (DH) | Replica-Exchange MD from sequence | 100 ns |
| sCaM-1 + DH | 2 MD refined sCaM-1 structures + DH docked at 4 sites | $2 \times 4 \times 3 \times 1.2 \mu s = 28.8 \mu s$ |
| sCaM-4 + DH | 2 MD refined sCaM-4 structures + DH docked at 4 sites | $2 \times 4 \times 3 \times 1.2 \mu s = 28.8 \mu s$ |
| mCaM + DH | Representative structure of MD simulations of 4Q5U + DH docked at 4 sites | $1 \times 4 \times 3 \times 1.2 \mu s = 14.4 \mu s$ |
| sCaM-1 + DH + AID | Representative sCaM-1 + DH at each site with AID fused to DH C-termini | $8 \times 1 \mu s = 8 \mu s$ |
| sCaM-4 + DH + AID | Representative sCaM-4 + DH at each site with AID fused to DH C-termini | $8 \times 1 \mu s = 8 \mu s$ |
| mCaM + DH + AID | Representative mCaM + DH at each site with AID fused to DH C-termini | $4 \times 1 \mu s = 4 \mu s$ |
| | | total: $\sim 110 \mu s$ |

**Figure S1:**
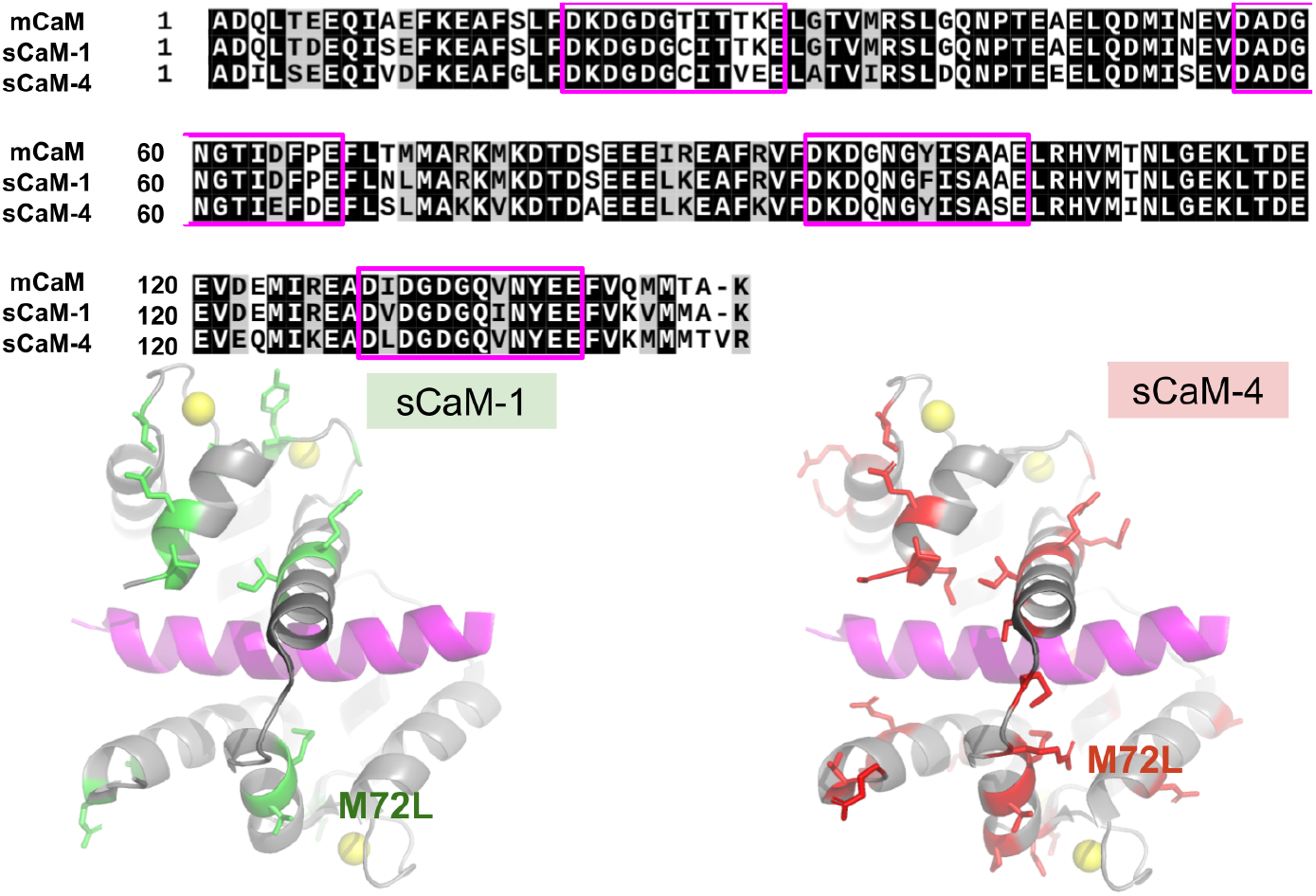
Sequence comparison between mammalian CaM (mCaM) and the sCaM-1/sCaM-4 soybean isoforms. The EF-hand residues are highlighted with magenta boxes. The lower panel highlights residues in soybean CaMs that are conserved in the human CaM/CaMBR complex structure (PDB 4Q5U [27]). Unique residues are shown in sticks and colored green and red for sCaM-1 and sCaM-4, respectively.

## S2 Supplement Method

### S2.1 Protocol and flags of Rosetta Comparative Modeling

#### Listing 1: HybridizeMover Protocal

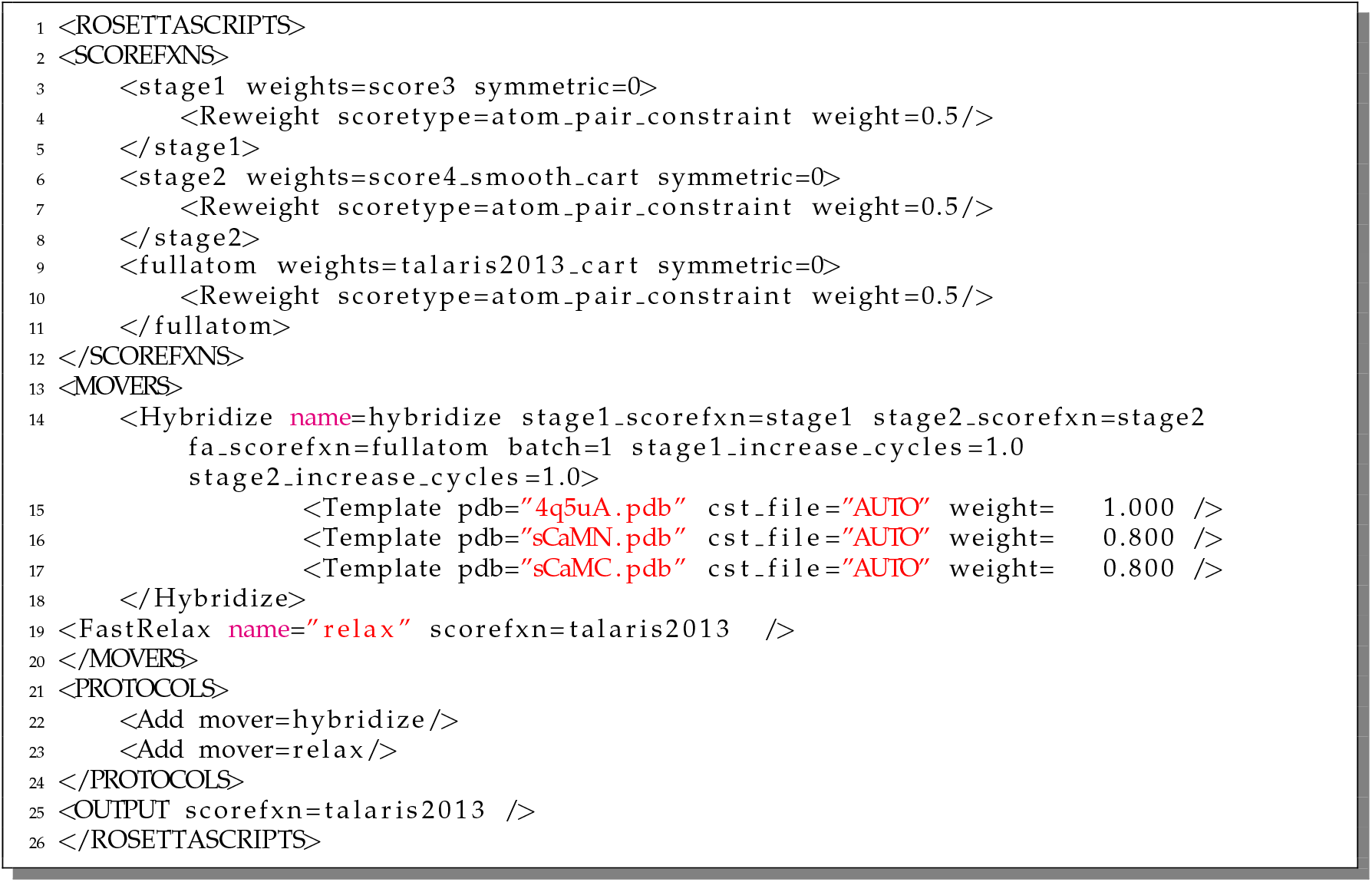

#### Flags

##### Listing 2: Flags

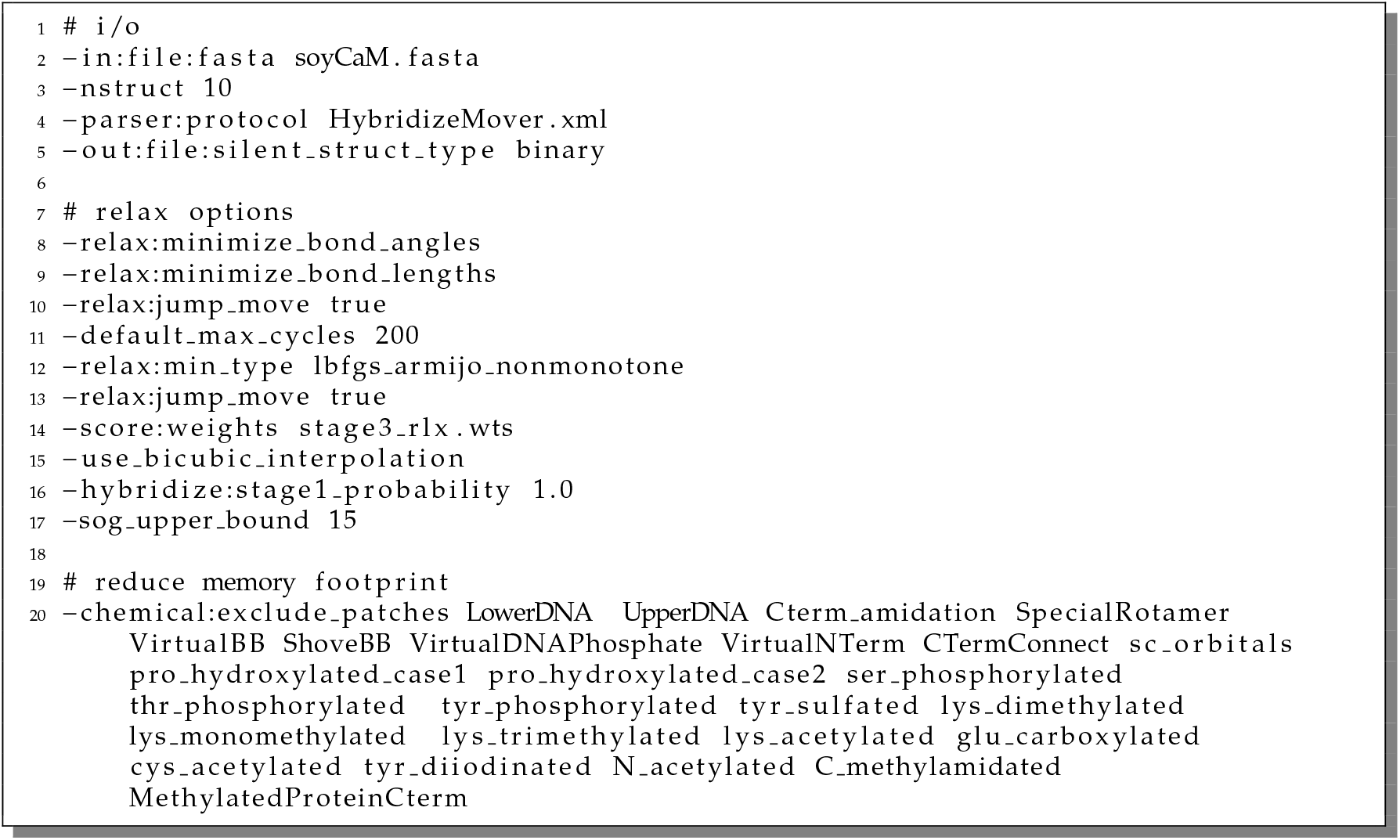

**Figure S2:**
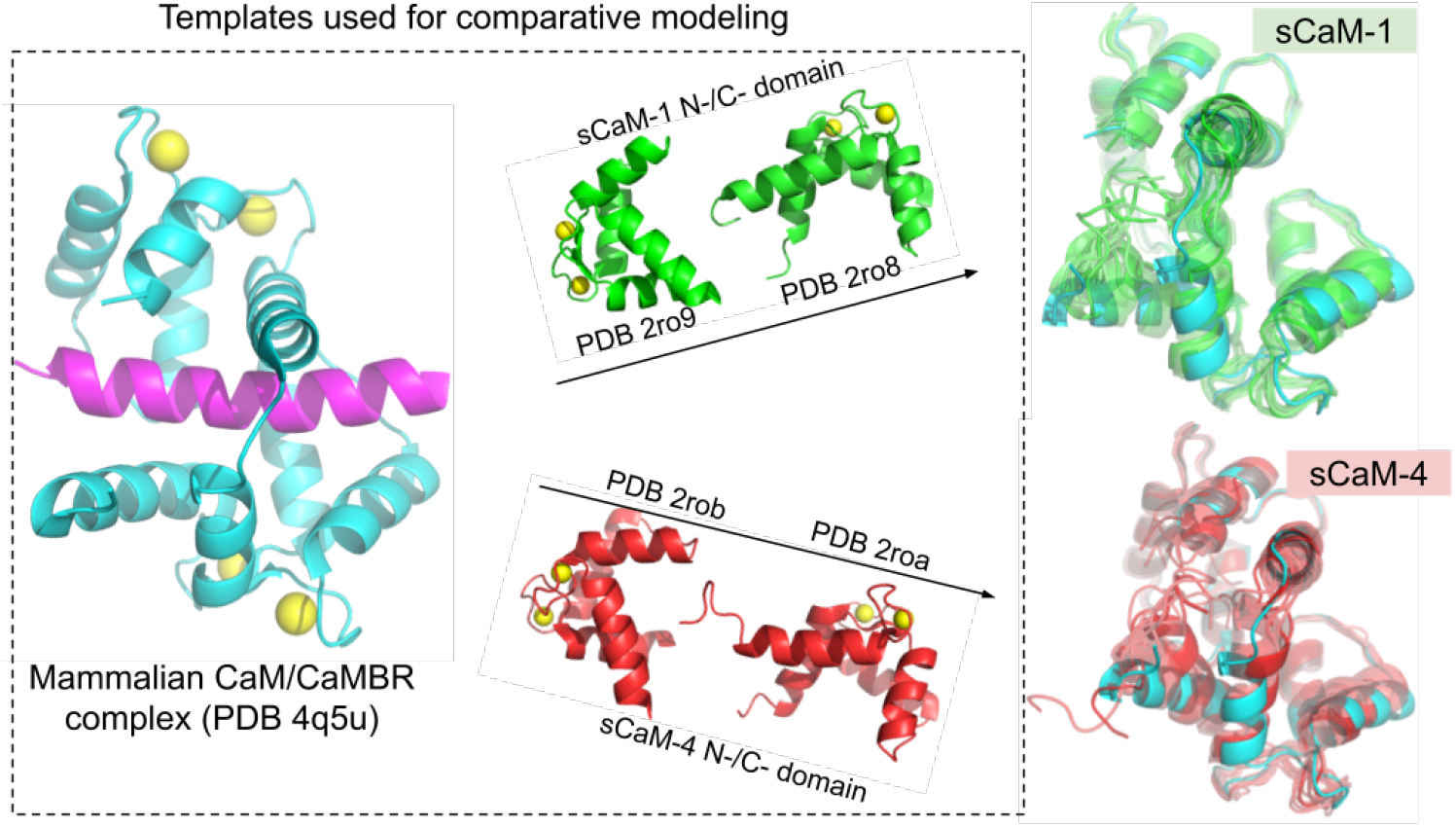
Rosetta comparative modeling of sCaM/CaMBR complex structures using human CaM/CaMBR crystal structure (PDB ID: 4q5u) and isolated sCaM domain structures as templates. The 10 modelled structures are superimposed on mammalian CaM template and are color green and red for sCaM-1 and sCaM-4, respectively.

### S2.2 Homology modeling combined with MD refinement gives reliable sCaM/-CaMBR complex structures

Although the separate N-/C-domains structures of sCaM-1 and sCaM-4 with Ca^2+^bound in the EF-hands have been resolved [11], complete sCaM-1/4 structures complexed with CaMBR of CaN are not yet unavailable. We thus utilized the Rosetta comparative modeling tools [32] to build the complete structures of sCaM isofroms to provide a structural basis for exploring CaM/-CaN interactions. Starting from the highest-scored Rosetta modeled structures, we performed 2 *μ*s MD in triplicates to refine the modeled structures as well as to investigate the structural and dynamic features of the sCaM isoforms. In addition, we also performed MD for the human CaM/CaMBR complex starting from the crystal structure PDB 4Q5U as a comparison.

**Figure S3:**
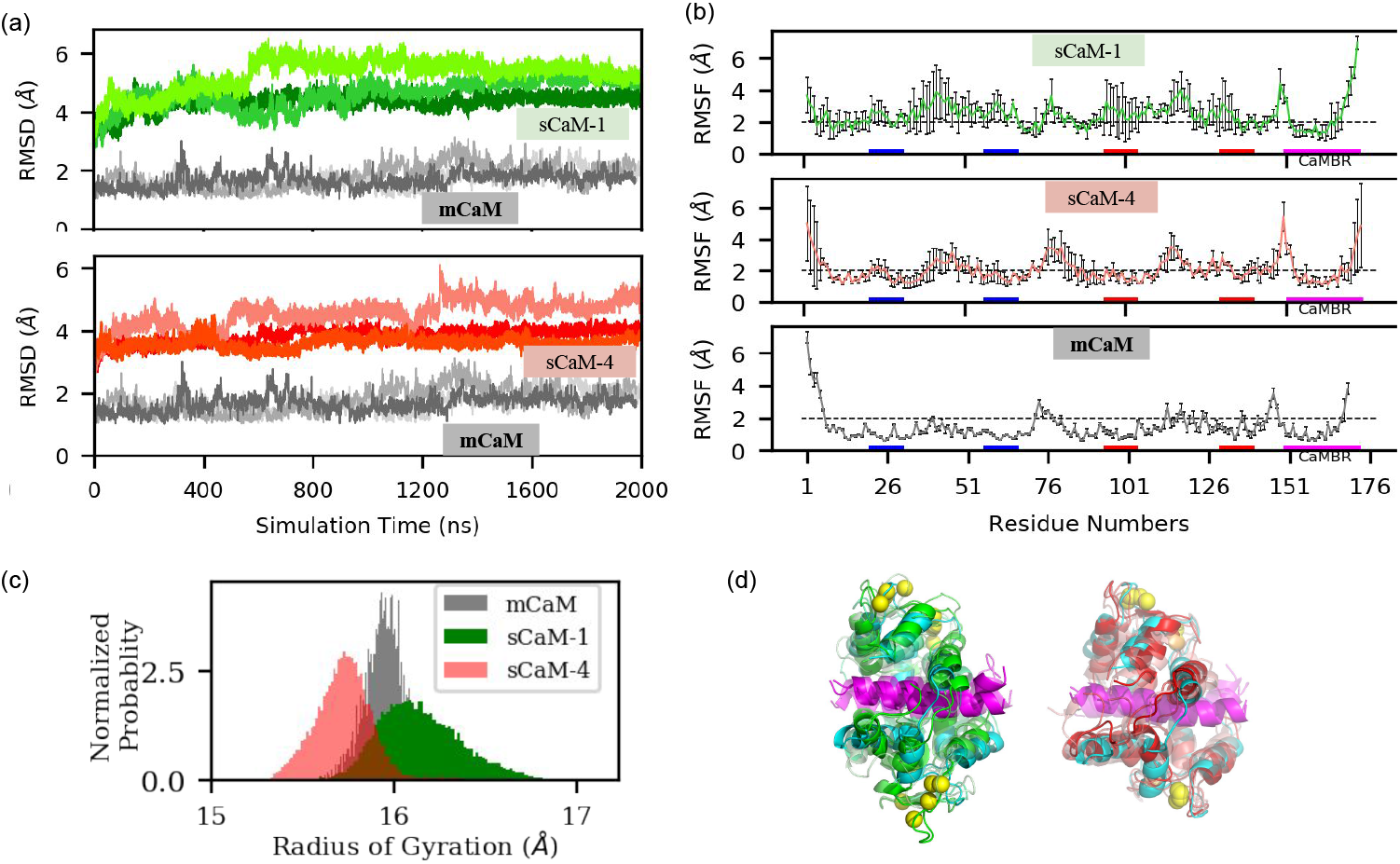
(a) root mean squared deviations (RMSD) of backbone atom of CaM/CaMBR. The first frame of the trajectory was used as reference of RMSD calculation. (b) root mean squared fluctuations (RMSF) of non-hydrogen atom in the triplicate MD simulations. The blue and red bars at the x axis represent the EF hands in the N and C domain, respectively. All frames were fitted to the first frame before RMSF calculation. The error bars are standard deviations calculated from the triplicate runs. (c) Radius of gyration distribution (d) Representative structures of sCaM-1 (green) and sCaM-4 (red) from triplicate MD simulations. The human CaM/CaMBR structure (PDB 4Q5U [27]) is colored cyan with CaMBR colored magenta. Ca^2+^ ions are represented as yellow balls.

As shown in Fig. S3a, the backbone atom root mean squared deviations (RMSD) values of sCaM-1 and sCaM-4 converges around 5 and 4 Å respectively. Meanwhile, the mCaM case started from the crystal structure maintains low RMSD values around 2 Å. The larger RMSD values for sCaM-1 and sCaM-4 are expected and it reflects the structural relaxation of the comparative modeled structures relative to its initial structure. Despite larger RMSD values for the two sCaM isoforms, these MD refinements suggest that modeled structures are stable over the 2 *μ*s time-scale simulations. More importantly, electrophoresis experiments of purified sCaMs using SDS-polyacrylamide gel in the presence of 5 mM CaCl_2_ [5] has shown that sCaM-4 has greater mobility than sCaM-1, implying that Ca^2+^bound sCaM-4 is more compact. Our MD simulations support this experimental observation, as shown in Fig. S3d, sCaM-1 has significantly larger radius of gyration than sCaM-4. Experiments also show that Ca^2+^ binding causes more hydrophobic patch exposure in sCaM-1 than sCaM-4 as measured by 1-anilino-8-naphthalenesulfonate (ANS) fluorescence spectroscopy [11]. The less-packed hydrophobic core of sCaM-1 might be the reason why it is more extended than sCaM-4. By integration of comparative modeling with appropriately selected templates and extensive atomistic MD simulation refinement, we were able to model reliable complete sCaM structures bound with CaMBR of CaN, as evident by 1) RMSD values that are converged over the *μ*s MD time-scale. Compact structure of sCaM-4 that is consistent with experimental observations. This integrated protocol has been widely used to build protein structures that have eluded experimental structural determination [66, 67].

**Figure S4:**
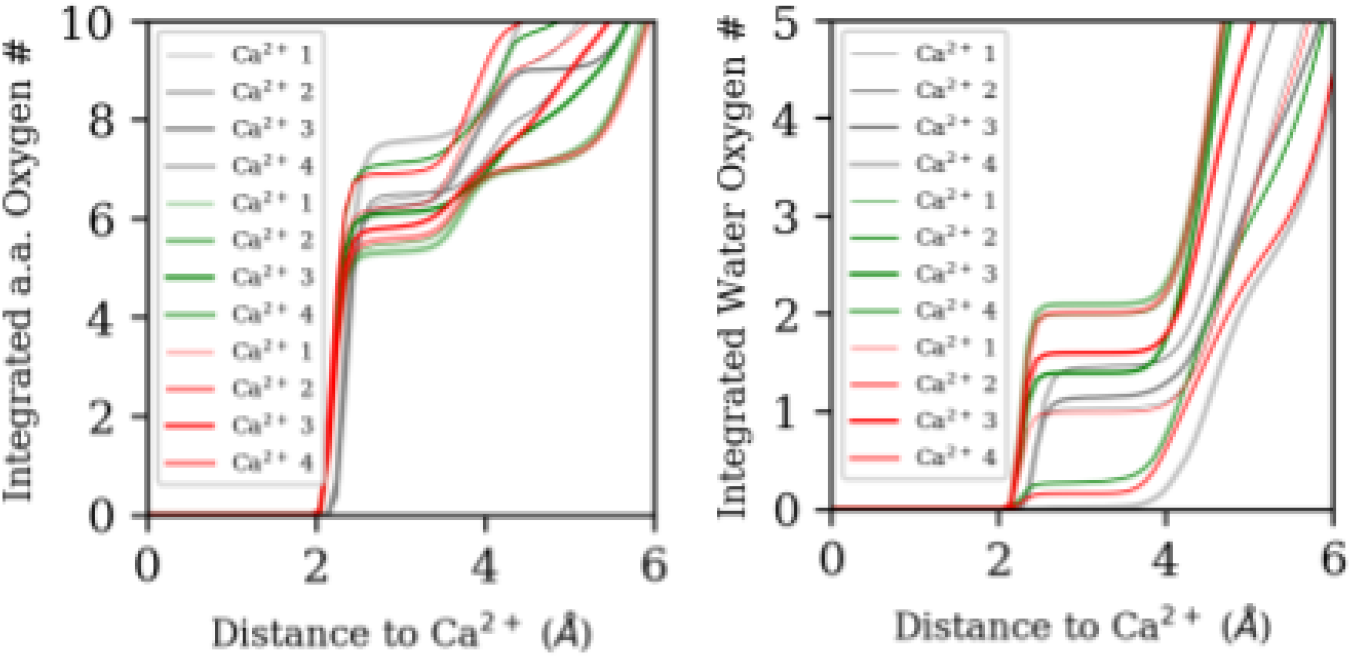
Integrated amino acid (a.a.,) and water oxygen numbers around Ca^2+^ ions.

### S2.3 Clustering analysis on sCaM MD trajectory to identify representative structures that will be used to dock with DH

To perform the clustering analysis, we first divided the triplicate MD runs for each sCaM into two groups, based on the RMSD values (Fig. S5). Thus for each sCaM, we have two groups A and B. For each group, a clustering analysis were performed using the cpptraj module of Amber with the RMS of backbone atoms as distance metric. The hierarchical agglomerative (bottom-up) approach was used and the MD frames were divided into ten clusters. The representative structure of the most populated cluster was picked out (the bottom panel of Fig. S5)

**Figure S5:**
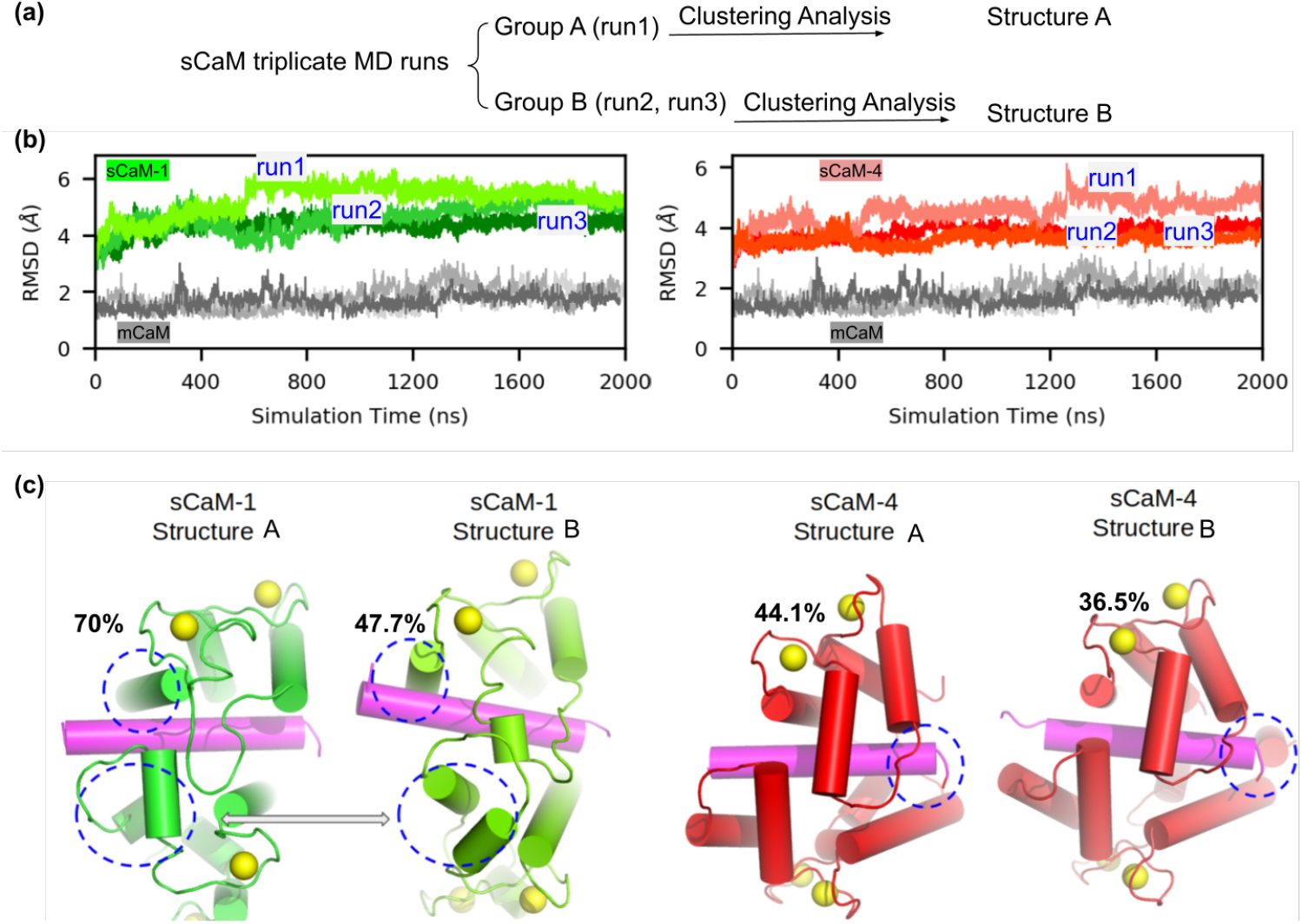
(a) Strategy of clustering analysis on the MD refinement trajectories. (b) Backbone atom RMSD of triplicate MD refinement simulations. The first frame was used as reference. (c) Two representative structures for each sCaM were selected to dock distal helix. The blue dashed circles highlight the structural difference of the two structures of the same sCaM.

### S2.4 REMD of distal helix and putative docking sites on CaM surface

**Figure S6:**
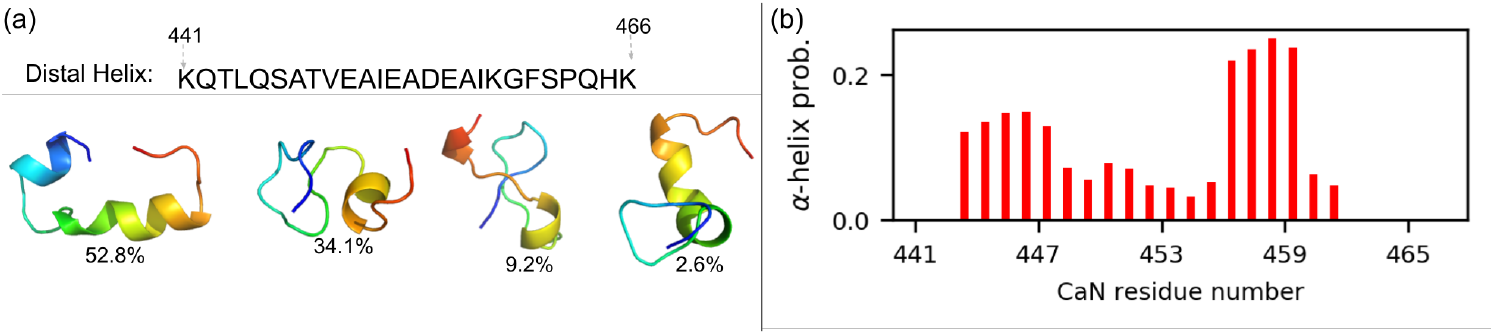
Structural property of distal helix sampled via 100 ns REMD simulation. (a) Sequence and representative structures of the first four most populated clusters. (b) *α*-helix probability of each residue.

**Table S2:** Residues at the each tentative binding site on sCaM-1/sCaM-4 used in ZDOCK to predict distal helix interaction at each site. The slightly different residues at each site between hCaM and sCaM-1/sCaM-4 was to make sure these selected residues are exposed on surface.

| Tentative Site | hCaM | sCaM-1 Residues | sCaM-4 Residues |
| --- | --- | --- | --- |
| Site A | R86, F89, V142, Y138 | K86, R90, Y138, V142 | K86, K90, Y138, K143 |
| Site B | R106, I125, D118, D122 | R106, D118, D122, R126 | R106, D118, E122, K126 |
| Site C | I9, F12, L69, F65 | I9, K13, F65, N70 | V10, K13, F65, S70 |
| Site D | K30 ,T34, R37, Q49 | K30 ,V35, S38, Q49 | E30 ,T34, R37, Q49 |

**Figure S7:**
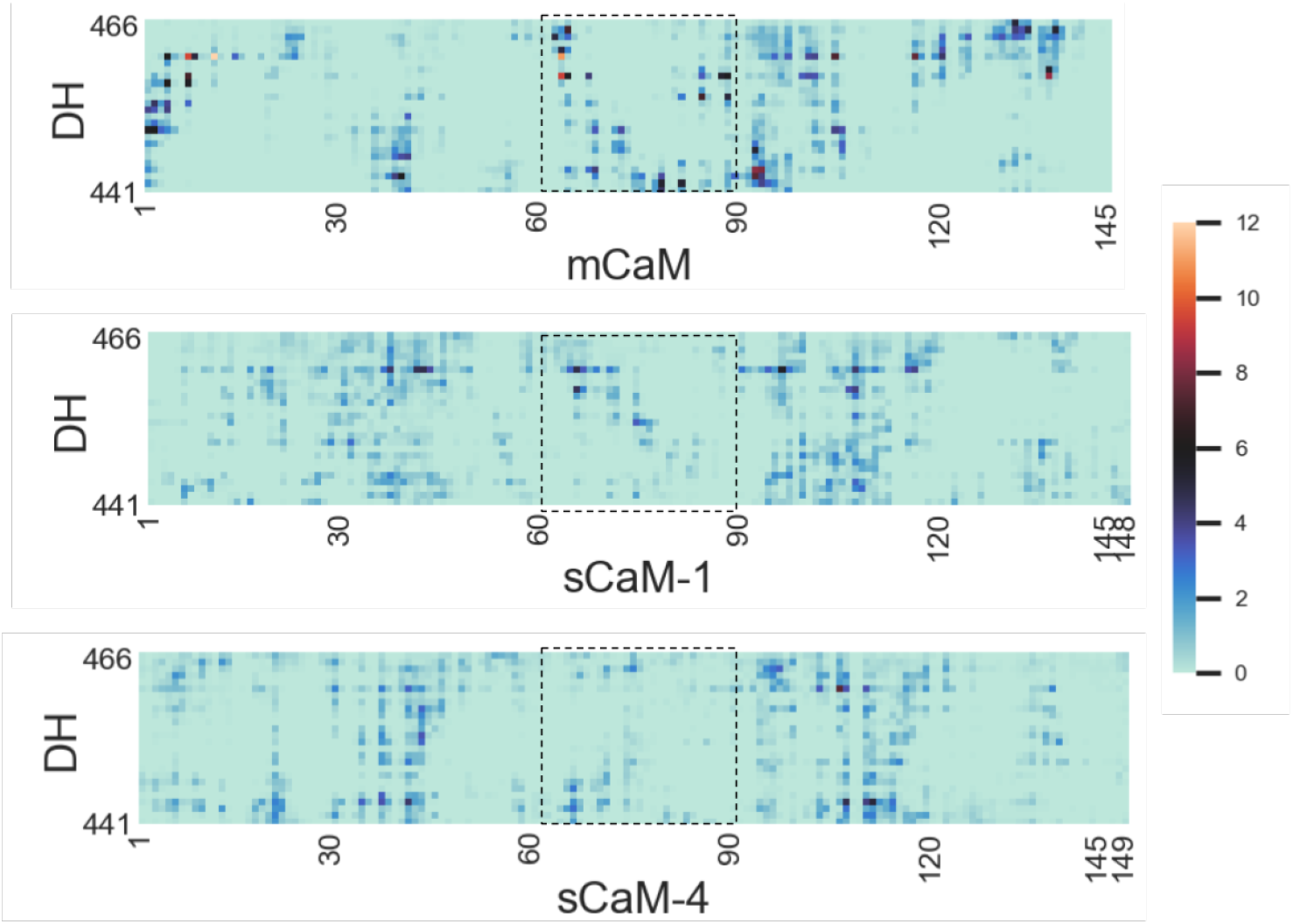
Contact map analysis between DH and CaM. The dashed boxes highlight the regions where sCaM-1 and mCaM have similar contact that differs from sCaM-4. The contact is defined by atoms that are within 7 Å. All contacts from the atoms comprising the residue are summed up to serve as reside-residue contact data shown here. Thus the color bar represents the total percentages of all atoms in one residue that have contacts with other residues

### S2.5 The functional consequence of the distal helix/CaM interaction is quantitatively explained by a tether model that gives [AID]_eff_ concentration as the indication of CaN activation

Previously we have successfully used *K_m_* to validate our simulations under the hypothesis that the binding of CaM to CaN RD alters the AID distribution around the active site, which can be shown in the apparent *K_m_* measured by experiments [30]. To relate the distal helix/CaM interaction to actual CaN activation, we utilized a tethering model our lab developed in [30] which gives the effective AID concentration ([AID]_eff_) near the catalytic site of CaN. Our hypothesis is that larger [AID]_eff_ leads to larger *K_D_* of substrate. Here we briefly summarize the key idea of the tethering model: in the absence of CaM, the regulatory domain of CaN is treated as a random coil and the AID is tethered by this coil (Fig. 1). The [AID]_eff_ around the catalytic site is determined by the length of the coil: when the length shortens, the [AID]_eff_ is reduced near the catalytic site, which reduces competition for substrate binding. Interacting with CaM will shorten the length of regulatory domain as the CaMBR and distal helix regions become rigid and locked by CaM, leading to much smaller [AID]_eff_. Quantitatively, the [AID]_eff_ is given by Van Valen et al [68],

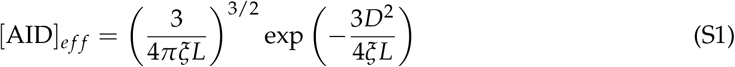

where *D* depicts the distance between N-termini of regulatory domain and catalytic site, which we assume is the distance between M387 and E481 in PDB 4OR9 [30].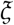 is the persistence length determined from literature [69]. *L* is linker length that we sampled from MD simulations. To obtain *L* values, we started from MD refined CaM/DH complex structures and added the AID fragment to the system and performed extensive all-atom MD to sample the relative position of AID around CaM as originally described in [30]. We measured the distance between the center of mass (COM) of AID to the COM of CaM (Fig. S8a,b) and used the values as *L* in Eq. S1. The distance distribution shows that the three CaMs have *L* values ranging from 18 to 40 Å.

Putting the values in Fig. S8b as *L* into Eq. S1, we obtained the [AID]_eff_ distribution for the three CaMs. The [AID]_eff_ distribution is much like the CaM-AID distance distribution. For interpretation, we assume the [AID]_eff_ value of the highest peak of mCaM is the critical value and the [AID]_eff_s that are smaller and larger than this critical value will exert activating and non-activating effect on CaN, respectively (the yellow line in Fig. S8c). We calculated the areas of the activating and non-activating peaks in Fig. S8c to assess the overall effect of these CaMs on CaN. By this measurement, we show that mCaM and sCaM-1 both have ~65 % activating effect on CaN while sCaM-4 only has 46 % (Fig. S8d). These trends suggest that sCaM-4’s weaker CaN activating power is due to its more non-activating effect on CaN than sCaM-1 and mCaM. In other words, all CaMs are capable of activating CaN, but sCaM-4 to a lesser extent.

**Figure S8:**
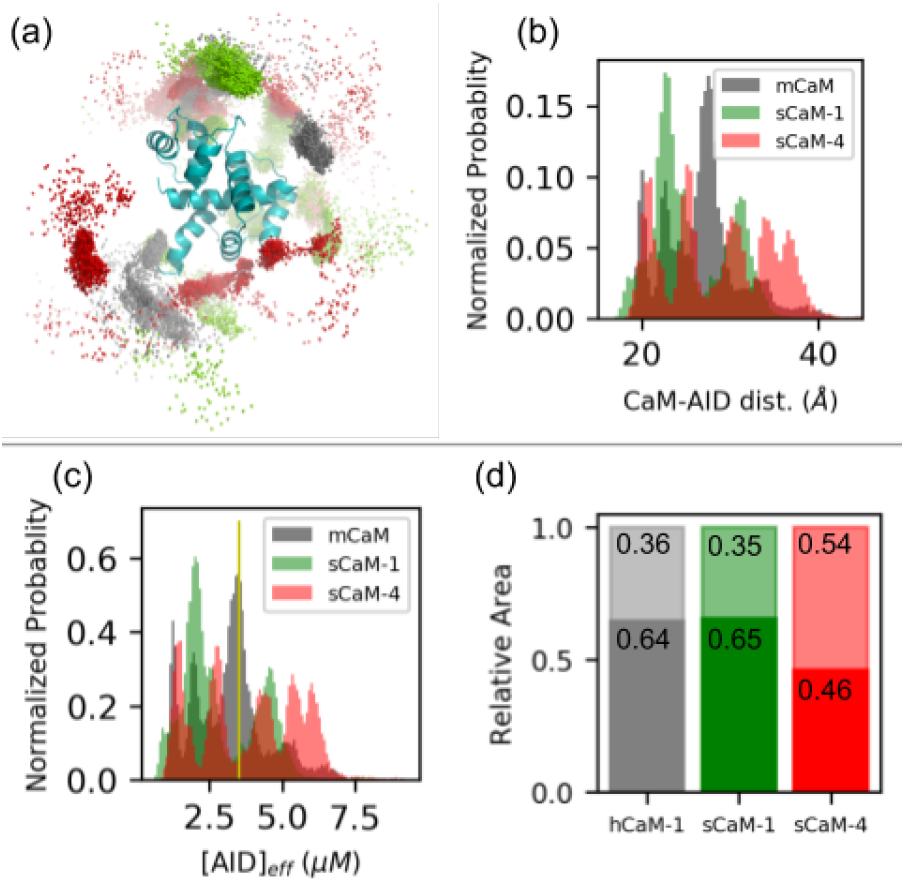
(a) Distribution of AID center of mass (COM) around CaM. (b) Distance between the COM of CaM and COM of AID. (c) The estimated effective AID concentration ([AID]_eff_) calculated via the tethering model with the CaM-AID distance as input. (d) Ratio of activating area versus inhibitory area calculated from panel c. Using [AID]_eff_ = 3.5*μ*M as thresholding value.

